# V-pipe 3.0: a sustainable pipeline for within-sample viral genetic diversity estimation

**DOI:** 10.1101/2023.10.16.562462

**Authors:** Lara Fuhrmann, Kim Philipp Jablonski, Ivan Topolsky, Aashil A Batavia, Nico Borgsmüller, Pelin Icer Baykal, Matteo Carrara, Chaoran Chen, Arthur Dondi, Monica Dragan, David Dreifuss, Anika John, Benjamin Langer, Michal Okoniewski, Louis du Plessis, Uwe Schmitt, Franziska Singer, Tanja Stadler, Niko Beerenwinkel

## Abstract

The large amount and diversity of viral genomic datasets generated by next-generation sequencing technologies poses a set of challenges for computational data analysis workflows, including rigorous quality control, adaptation to higher sample coverage, and tailored steps for specific applications. Here, we present V-pipe 3.0, a computational pipeline designed for analyzing next-generation sequencing data of short viral genomes. It is developed to enable reproducible, scalable, adaptable, and transparent inference of genetic diversity of viral samples. By presenting two large-scale data analysis projects, we demonstrate the effectiveness of V-pipe 3.0 in supporting sustainable viral genomic data science.

## 1 Background

With the advent of next-generation sequencing (NGS) technologies, large amounts of viral genomic data are being generated, which can no longer be easily analyzed on personal computers [1]. As this availability of high-coverage data sets brings interesting research opportunities but also computational challenges, many new processing and analysis tools are being developed. In particular, new possibilities of characterizing viral variants and analyzing the genetic diversity of viral sequencing samples have emerged [2, 3]. While inter-host variability describes how viral strains differ between separate hosts, within-host variability measures the diversity of viral strains within a single host. Within-host genetic diversity is thus especially relevant to understanding disease progression and treatment options [4, 5]. In addition to clinical or experimental samples, there has been an increasing abundance of environmental samples also showing within-sample variability, such as wastewater samples. These samples possess a diverse array of viruses, enabling the monitoring of pathogens on a larger scale, encompassing cities, regions, and countries [6, 7].

For estimation of within-sample diversity from NGS samples, several data processing steps and tools are needed. Due to the complexity of the data, these tools are usually executed as part of a processing workflow. Typically they combine tools for quality control, sequence alignment, consensus sequence assembly, diversity estimation, and result visualization. Various workflows have been proposed which try to accomplish these goals including V-pipe [8], ViralFlow [9], nf-core/viralrecon [10] and HAPHPIPE [11]. The adaptability of these workflows becomes crucial as different types of viruses require tailored analysis approaches. This need became evident during the SARS-CoV-2 pandemic, emphasizing the rapid emergence of specific requirements vital to public health [12]. For example, sequencing samples originating from diverse sources, such as clinical or wastewater settings, require application-specific processing steps that need to be supported in the same workflow.

Another effect of the SARS-CoV-2 pandemic is that a substantial increase in sequencing capacities has led to unprecedentedly large numbers of samples becoming publicly available, e.g., on the European Nucleotide Archive (ENA; [13]) or GenBank [14]. Analysis workflows need to be able to handle such large amounts of data in order to be beneficial to public health and epidemiological advances. Hence, it is critical for workflows to not only include a broad range of functionalities, but also to promote sustainable data processing practices to ensure their effectiveness and long-term success.

NGS data processing workflows offer a range of diversity estimation approaches at different spatial genomic scales: mutation calling, local and global haplotype. Mutation calling refers to detecting genetic mutations or variations at specific positions within the genome. Global haplotypes refer to the reconstruction of complete haplotypes that span the entire length of the viral genome. On the other hand, local haplotypes focus on identifying mutations within a single read. The reconstruction of global haplotypes is more complicated as multiple reads need to be assembled together to cover a whole genome, but it provides a more comprehensive measure of viral diversity [15].

As the methodologies for viral diversity estimation and data sources can be heterogeneous, understanding the performance of each tool and benchmarking them in a realistic way is difficult. Additionally, different methods may excel in different scenarios. Therefore, continuous benchmarking of these methods is crucial to identify the most suitable one for a given data source and scenario. Consequently, it is important to provide data analysis procedures as publicly available workflows designed in a sustainable manner. This approach facilitates continuous re-evaluation of the benchmarking workflow with new and updated parameter settings. This is needed as new methods are being developed which have to be compared to already existing ones, new test data sets become available, either new synthetic data sets with new simulation setups, or real data sets with new experimental setups. Finally, completely new application domains can appear which requires adapting the existing benchmarking workflow.

Here, we present V-pipe 3.0, a sustainable data analysis workflow for diversity estimation from viral NGS samples. Sustainability comprises reproducibility, scalability, adaptability and transparency of the workflow [16]. V-pipe 3.0 builds upon the foundation of V-pipe [8], but has undergone significant extensions and refinements to address new challenges and adhere to sustainable data processing standards [16]. We highlight how the workflow has been designed to achieve these properties and describe how they have been crucial for the application of V-pipe 3.0 to large-scale data analysis projects. In particular, we present a new and efficient workflow that enables the processing of hundreds of thousands of samples. We demonstrate how automated source code testing makes it possible to quickly make new functionalities and bug fixes available to end users and how its modular design allows to quickly implement application-specific features. Further, for the evaluation of suitable genetic viral diversity estimation, we added a benchmarking module. This module itself is sustainably implemented and it enables adding new methods and test data sets. We demonstrate its use by conducting a benchmarking study where we apply a set of global haplotype reconstruction methods to both synthetic and real data sets. Lastly, we compare V-pipe 3.0 to workflows for similar applications, provide an overview of their functionalities, and compare their structures in terms of sustainability.

V-pipe 3.0 is publicly available on GitHub [17].

## 2 Results

V-pipe 3.0 is a bioinformatics workflow which combines various tools for analyzing viral NGS data (Table 1). V-pipe 3.0 is based on V-pipe, a pipeline designed for analyzing NGS data of short viral genomes [8] and extends it not only in terms of functionalities but also by consistently implementing principles of sustainable data analysis. In the initial step of the pipeline, the raw sequencing reads in fastq format undergo a quality control process. Following this, the reads are aligned, and subsequently, the user-specified diversity estimation methods are executed (Figure 1). To ensure sustainable data analysis using V-pipe 3.0, we followed the hierarchy of sustainability proposed in [16] and created a reproducible, scalable, adaptable, and transparent workflow. It has been widely recognized that these aspects are crucial to scientific progress but often lacking in current literature [34, 35]. In the following, we will provide a detailed explanation of the reimplementation and extensions that were undertaken during the development of V-pipe 3.0. To demonstrate that V-pipe 3.0 effectively addresses the challenges of sustainable data analysis we follow the four aspects in Mölder’s hierarchy [16].

**Fig. 1:**
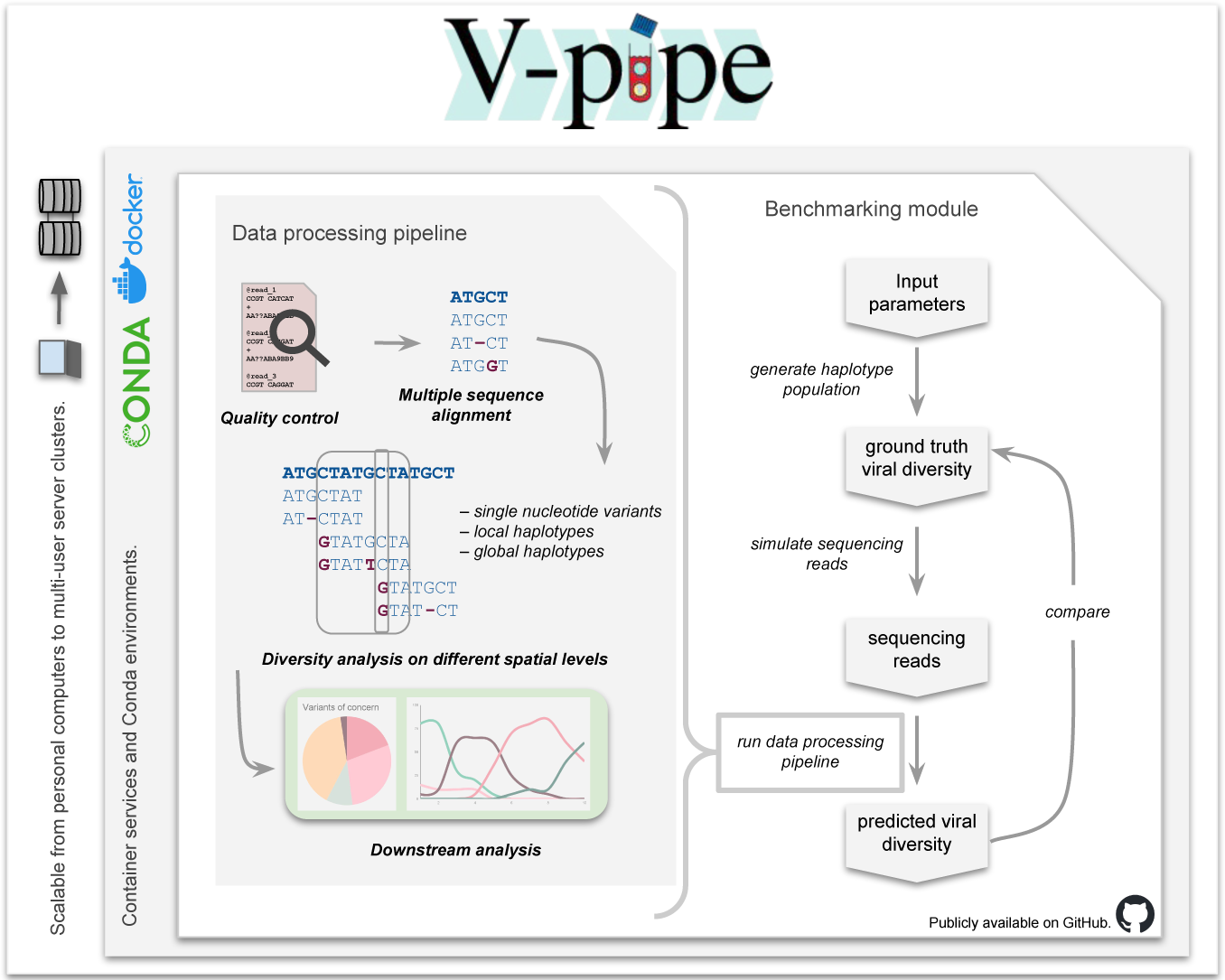
V-pipe 3.0 workflow overview. The data processing pipeline (left) provides four main steps: (1) Preprocessing of the raw reads including quality control, (2) multiple sequence alignment, (3) estimation of viral diversity by SNV, local and global haplotype calling, and (4) if applicable, downstream analysis. The V-pipe 3.0 benchmarking module (right) supports the evaluation of viral diversity estimation methods on simulated data and on real experimental data where the ground truth diversity is known by the experimental design. For the simulated samples, first, ground truth haplotype populations are generated and based on those, sequencing reads are simulated. Then, the simulated and real samples are processed by the methods in the study, and last, the predicted viral diversity is compared to the ground truth viral diversity using different metrics for example precision, recall, f1 and N50 score. V-pipe 3.0 is designed to facilitate efficient processing on personal computers as well as on computing clusters. V-pipe 3.0 automatically sets up the necessary Conda environments, installs all dependencies, and initializes the project structure. It is also accessible through a Docker container, which includes all software dependencies.

**Table 1:** Methods and tools per data processing step that are integrated in V-Pipe 3.0.

| Data processing task | Tool | Reference |
| --- | --- | --- |
| Quality control | PRINSEQ | [18] |
|  | FastQC | [19] |
| De novo assembly | VICUNA | [20] |
| Primer trimming | IVar | [21] |
|  | SAMtools | [22] |
| Aligner | BWA MEM | [23] |
|  | Bowtie 2 | [24] |
|  | minimap2 | [25] |
|  | ngshmmalign | [8] |
| Consensus sequence generation | SmallGenomeUtilites | [8] |
|  | BCFtools | [22, 26] |
| Mutation calling | LoFreq | [27] |
|  | ShoRAH | [28] |
| Local haplotype reconstruction | ShoRAH | [28] |
| Global haplotype reconstruction | PredictHaplo | [29] |
|  | HaploConduct | [30] |
|  | HaploClique | [31] |
|  | QuasiRecomb | [32] |
| SARS-CoV-2 wastewater surveillance | COJAC | [6] |
|  | LolliPop | [33] |

### 2.1 Reproducibility

Reproducibility allows other researchers to execute an existing workflow and obtain the exact same results as the original workflow authors. To achieve this goal, we define all software dependencies in Conda environments which makes V-pipe 3.0 portable between different computing platforms. That way, V-pipe 3.0 can be executed without complicated, manual installation procedures. To ensure successful installation and reproducible execution on different systems, we use GitHub Actions [36] for automatic test installations on Mac OS and Linux systems, and for end-to-end tests by executing tutorials with example data.

The reproducibility of V-pipe 3.0 results is strongly dependent on the reproducibility of the integrated methods. One core functionality of V-pipe 3.0 is the estimation of viral genetic diversity. A multitude of viral diversity estimation tools exist, making it challenging for users to determine the appropriate tool for their samples. Additionally, the choice of method depends on the desired downstream analysis of the results. Therefore, we created a Snakemake based workflow as part of V-pipe 3.0 which automatically applies a set of selected tools to various synthetic and real data sets, computes their respective performances in terms of precision and recall, and summarizes the results.

The benchmarking workflow is itself sustainably implemented. Adding new tools and data sets to this benchmark is very easy and only requires the addition of a single file and no further modifications of the workflow. By incorporating the benchmarking module, we enhance the sustainability of V-pipe 3.0, as robust and continuous benchmarking all integrated software components makes the workflow more adaptable to new data sets and its results reliable. Moreover, this framework facilitates the easy assessment of new diversity estimation methods enabling extensions of V-pipe 3.0 to be implemented in a reproducible fashion. As a concrete demonstration of the benchmarking module’s effectiveness, we conducted a benchmarking study focused on global haplotype reconstruction (Section 3.3).

### 2.2 Scalability

Scalability allows the workflow to handle and process increasing amounts of data without compromising on performance or efficiency. To achieve scalability, we utilize efficient programming techniques to execute jobs on a computing cluster, ensuring optimal performance. For example, we dynamically specify cluster resources to adapt to the specific data requirements, facilitating smoother deployment on new cluster environments and enable the parallel execution of unrelated data analysis steps. Furthermore, we validate user configuration files using JSON Schema [37] during startup to identify potential runtime errors early. Lastly, we split centralized tasks among multiple compute nodes and perform per-sample distributed computation of summary statistics. In order to make large-scale analyses of public data sets easier, V-pipe 3.0 includes an input data retrieval functionality which requires a set of SRA accession numbers [13] as input and automatically downloads all data files needed to run the whole workflow. Further, scripts are available which facilitate the unattended mass-import of raw files as produced by Illumina’s demultiplexing software into the structure that V-pipe 3.0 expects as input. To help with common post-processing steps, we have added scripts to facilitate the SRA and GISAID database upload of compressed raw reads and of generated consensus sequences, including the summary quality reports assessing the plausibility of frameshift-causing insertions and deletions. With these features, V-pipe 3.0 has been shown to handle more than 100,000 samples efficiently [38–41].

### 2.3 Adaptability

Adaptability refers to making it easy for other researchers to build upon an existing workflow and extend it for their application- and domain-specific needs. To ensure that new functionalities can be quickly added to the workflow without compromising correctness, we track the development using git and run automated integration and unit tests using GitHub Actions workflows [36] on every commit submitted to the repository. We use data sets from different viruses in our tests to make sure that V-pipe 3.0 and the newly added features are running successfully from start to end.

To demonstrate the ease with which new software components and scrips can be introduced we added two methods for viral diversity estimation: first, PredictHaplo [29] a well-performing global haplotype reconstruction method, and second, a script for the computation of within-sample diversity indices [42], like Shannon Entropy or population nucleotide diversity. The indices are often applied to compare diversity between samples and have been used for the estimation of time since infection [43]. The addition of new methods requires only the definition of a Conda environment with the required software dependencies and the definition of a Snakemake rule executing the method or script. This ensures that new functionalities are easily integrated into V-pipe 3.0.

Further, V-pipe 3.0 can be easily optimized for different viruses through its configuration setup. The base configuration is virus-agnostic while virus-specific settings (specific reference sequences, different alignment tools, etc.) can be easily plugged in. This allows a quick adaptation of V-pipe 3.0 to any virus, without requiring complex workflow changes. For example, we provide HIV- and SARS-CoV-2-specific configuration setups, which select appropriate reference files, read alignment software and post-processing steps. To show how to write such configuration files for other viruses, we added a monkeypox-specific configuration file (Figure 2). The configuration defines which alignment and diversity estimation method should be applied, which reference should be used, and which outputs and processing steps should be run. Further, for each method, users can specify the parameter choices.

**Fig. 2:**
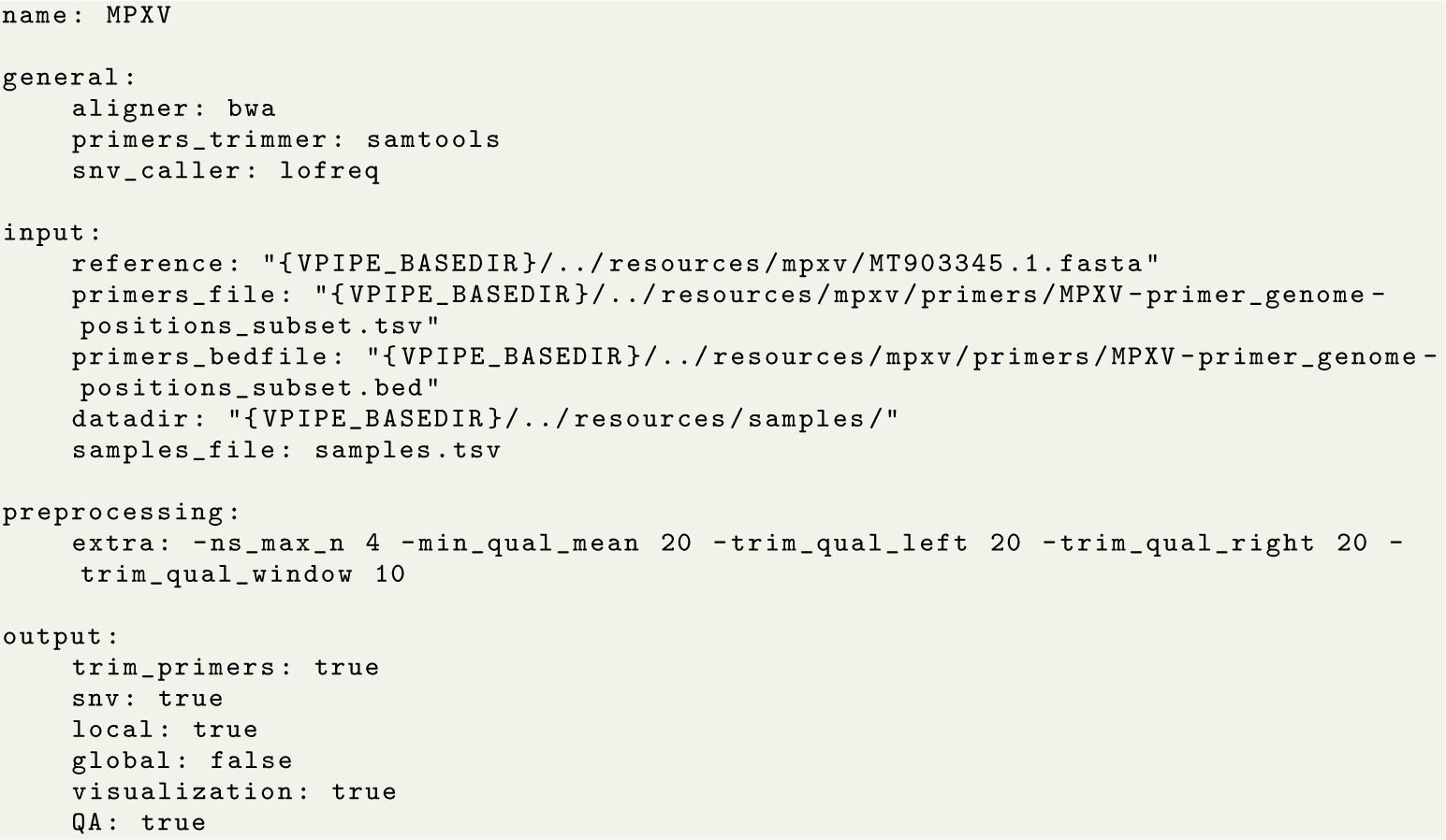
Example configuration file for monkeypox virus. User-specified aligner, primer trimming method, and the method for the diversity estimation are defined in the general section. Input like reference genome, primer file, and the directory of the samples are specified in the input section. In section preprocessing, extra command line parameters are passed to the preprocessing step. In section output, users can define their desired output of the pipeline. This example configuration file is available on GitHub [44].

### 2.4 Transparency

Transparency refers to the ability to easily comprehend a given workflow. This is particularly crucial for ensuring interpretability and facilitating efficient collaboration in large-scale projects with many stakeholders. V-pipe 3.0’s documentation is written as dynamic scripts which allows testing of the configuration options in an automated fashion and making sure they always represent the latest release version and do not contain outdated information. Additionally, V-pipe 3.0 offers a range of tutorials that cover various applications, including the processing of SARS-CoV-2 or HIV samples, as well as a tutorial specifically designed for processing wastewater samples.

In order to facilitate prompt user access to new functionalities and accelerate the onboarding process for new users, we provide four deployment methods: (1) a Bash script which automatically creates the required Conda environments, installs all dependencies and initializes a project structure, (2) the ability to use Snakemake’s snakedeploy tool to install V-pipe 3.0 in the standardized Snakemake fashion, (3) a Docker container [45] which is automatically generated for every new release and for the master branch of the git repository, and (4) the execution within a workflow execution service (WES), such as Sapporo [46], by fetching V-pipe from a tools repository service (TRS) such as WorkflowHub [47]. Further, V-pipe 3.0’s configuration definition summarizes the steps of the workflow in one single file and hence also facilitates information sharing between collaborators.

## 3 Applications

In the following, we present how sustainable data processing using V-pipe 3.0 was key to the successful execution of two large-scale national SARS-CoV-2 surveillance projects, and we demonstrate the benchmarking module by conducting a global haplotype reconstruction benchmarking study.

### 3.1 Swiss SARS-CoV-2 Sequencing Consortium

In the scope of the Swiss SARS-CoV-2 Sequencing Consortium [48], V-pipe 3.0 was consistently utilized to process sequencing data and generate consensus sequences. This continuous usage began with the first consortium sequencing run on 23 April 2020, and concluded when the consortium was dissolved in January 2023. V-pipe 3.0 demonstrated its adaptability by transitioning from its original focus on HIV to processing samples from SARS-CoV-2. The first Swiss SARS-CoV-2 case was reported on 25 February 2020 [49], and we submitted the first sequence processed by V-pipe 3.0 to GISAID on May 25th 2020 (accession number: EPI ISL 451681, sampled on 12th March 2020). The fast development and changing demands in the SARS-CoV-2 pandemic required the rapid development of new tools that had to be integrated in the processing pipeline, for example, the frameshift insertion/deletion checks as mentioned before. Apart from adaptability, portability and reproducibility were essential for this project, as it involved analysis conducted by different individuals from various academic groups on their own computing facilities. Since the consensus sequences and their Pango lineage [50] designations were reported to the Swiss Federal Office of Public Health to inform public health decision-making, reproducibility was essential to guarantee reliable, consistent, and trustworthy results. Further, V-pipe 3.0’s scalability to maximize the use of computational resources made it possible to handle the large amounts of clinical SARS-CoV-2 samples throughout the pandemic [51], which resulted in 74,409 consensus sequences being submitted to GISAID [52] as of 21-09-2023 (accessed 21-09-2023). At the peak of our efforts, V-pipe 3.0 processed up to 1500 clinical samples on a weekly basis (Figure 3A), providing a substantial part to the national surveillance efforts of circulating SARS-CoV-2 variants in Switzerland [38–40].

**Fig. 3:**
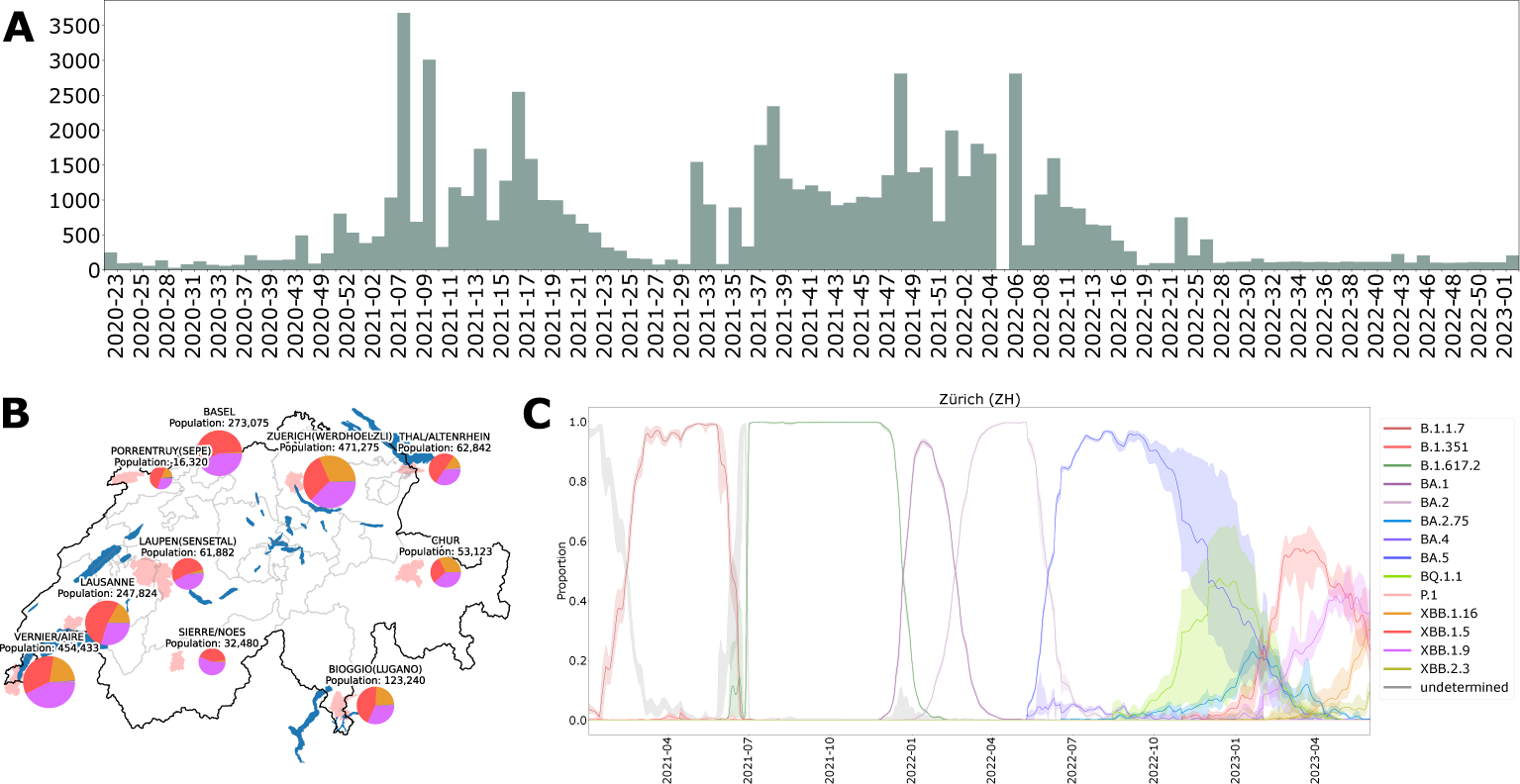
Swiss surveillance of SARS-CoV-2 genomic variants using V-Pipe 3.0. **A)** Number of weekly submission of SARS-CoV-2 consensus sequences from clinical samples to GISAID. Samples were processed with V-pipe 3.0. **B)** V-pipe 3.0’s surveillance of SARS-CoV-2 variants in wastewater samples from ten locations in Switzerland with relative abundances of variants. **C)** Time-series of relative variant abundances with 95% confidence bands of wastewater samples from Zurich using V-pipe 3.0.

### 3.2 Swiss surveillance of SARS-CoV-2 genomic variants in wastewater

Another successful application of V-pipe 3.0 has been the Swiss surveillance of SARS-CoV-2 genomic variants in wastewater [53] (Figure 3C). This category of samples contains mixtures of multiple SARS-CoV-2 lineages and workflows targeting diversity analysis are prime candidates for handling them. V-pipe 3.0 was used to analyze the sequencing data and to estimate the abundances of the circulating SARS-CoV-2 variants in Switzerland. In particular, the wastewater analysis enabled the early detection of new variants of concern such as Alpha (B.1.1.7) [6]. Starting in December 2020, V-pipe 3.0 has been continuously used to process wastewater samples from 6-10 different locations 3-7 times per week [53] (Figure 3B). Since then, V-pipe 3.0 has been the core of the automated monitoring of the circulating SARS-CoV-2 genomic variants in Swizterland (Figure 3C). The first 1823 out of the more than 6000 samples have already been submitted to the ENA project (PRJEB44932).

The complexity of the SARS-CoV-2 variant mixtures in wastewater samples required additions to the standard workflow, namely primer trimming and the newly developed methods COJAC [6] and LolliPop [33] for variant detection and time-series deconvolution of the variant mixtures. The modular and standard Snakemake structure of V-pipe 3.0 facilitated the integration of the new functionalities through adding new Snakemake rules for their execution. Lastly, the involvement of the large number of stakeholders and collaborators in the surveillance consortium of SARS-CoV-2 genomic variants in wastewater required transparency of the whole analysis pipeline. All stakeholders and developers had to be aware of the functionalities and steps of the data processing. This was possible through the modular structure and the clear configuration files used by V-pipe 3.0, as well as the fact that all parts of the pipeline are open source and their configuration automatically documented.

### 3.3 Global haplotype reconstruction benchmark

To showcase the strengths of V-pipe 3.0’s benchmarking module, we designed a global haplotype reconstruction benchmark study. Global haplotype reconstruction is a useful methodology in genetic research as it allows for a comprehensive understanding of the underlying genetic variations within a population. Due to the computational challenges involved in global haplotype reconstruction [54], it serves as a valuable application for the benchmarking module. Additionally, this benchmarking study provides an opportunity to evaluate new methods that could potentially be included in V-pipe 3.0. In our study, we compared the performance of the probabilistic method PredictHaplo and the graph-based methods CliqueSNV, HaploConduct, and HaploClique. We setup the benchmarking such that the methods were tested on two synthetic data sets and on one real data set.

Using the integrated synthetic data generation component of the module, we consider a genome of length 10, 000 bp, generate a population of 10 haplotypes (Population 1) and simulate Illumina reads of length 200 (Section 7.2). We vary the coverage between 500, 1000, 5000, 10, 000 in order to investigate how well the methods are able to recover low-frequency haplotypes as the coverage decreases.

We observe that PredictHaplo achieves perfect precision of 1 in all cases, CliqueSNV’s mean precision is between 0.60 and 0.68 with a slight increase with higher coverage (Figure 4A). In terms of recall, CliqueSNV features the highest recall of 0.5 *−* 0.6 which remains constant over all coverage values, while PredictHaplo’s recall increases up to 0.30 for the highest coverage of 10, 000. Consequently, the recall performance of CliqueSNV is less dependent on the coverage level when compared to PredictHaplo. Across all coverage values, CliqueSNV and PredictHaplo consistently achieve N50 scores of 10, 000, covering the entire genome length. In contrast, both HaploClique and HaploConduct fail to cover even a quarter of the genome, and show a precision and recall of 0 in all cases. This indicates that all sequences predicted by HaploClique and HaploConduct have relative edit distance greater than 0.01 to any true haplotype, and no true haplotypes are recovered. The poor performance could be attributed to HaploClique being executed with restricted clique size and maximal clique size, which may not be adequate for the assembly of longer regions. This parameter choice was necessary to prevent excessively long runtime and memory consumption. For all methods, we see a general trend of growing runtime with increasing coverage. CliqueSNV consistently requires the least amount of time to run, while PredictHaplo needs over an hour for the highest coverage (Figure 4A).

**Fig. 4:**
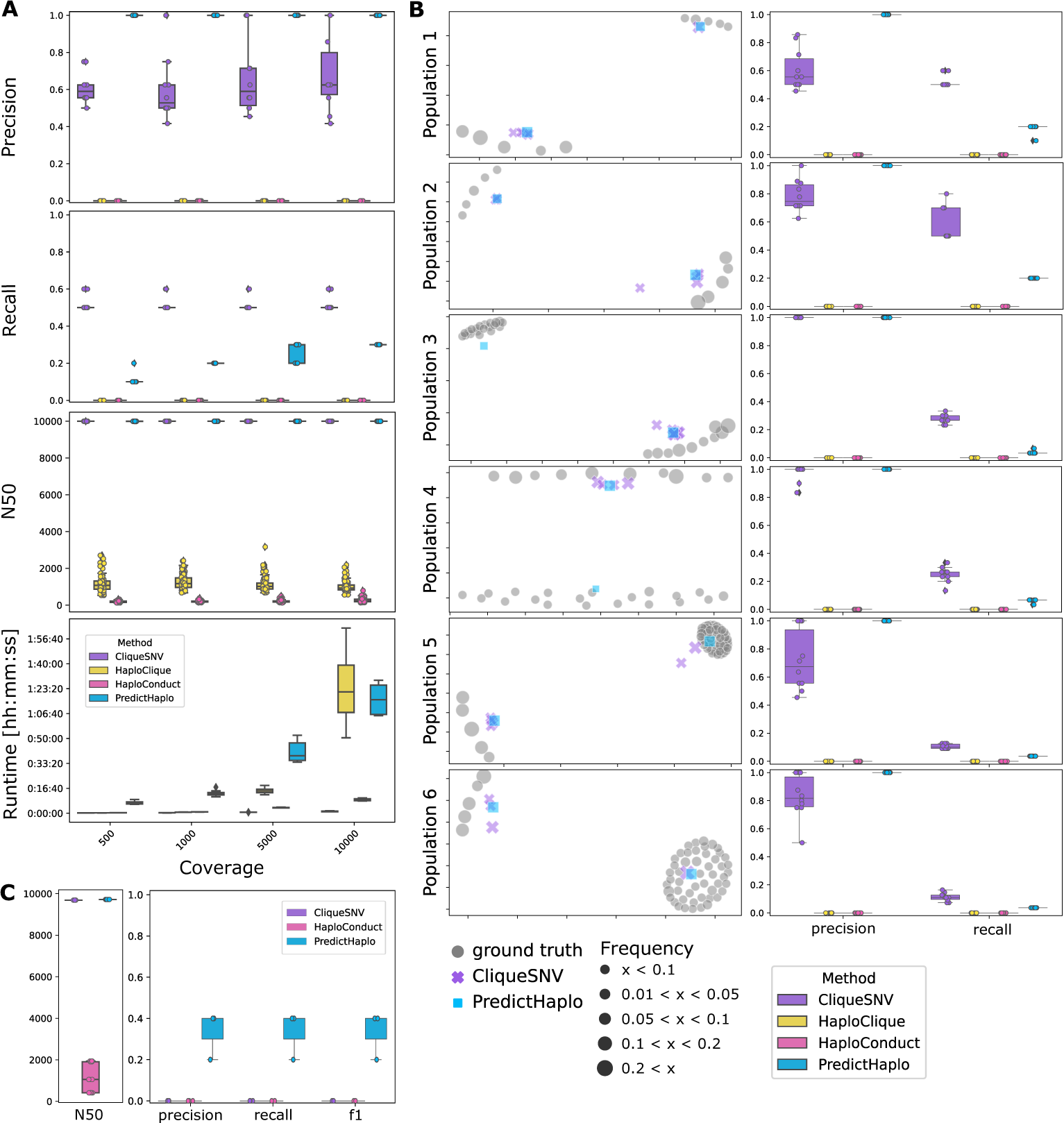
Benchmarking study for global haplotype reconstruction methods. **A)** Precision, recall, N50 score and runtime for simulated samples of varying coverage of population 1. **B)** Left: MDS plots of one example simulation replicate per haplotype population. Each point represents a sequence. Symbol size corresponds to the frequency of the respective haplotype in the sample. HaploClique and HaploConduct were excluded due to their poor performance. Right: Precision and recall plots for each haplotype population. Each marker represents one replicate sample. **C)** N50, precision, recall and f1 for PredictHaplo, CliqueSNV and HaploConduct on a real HIV-5-virus mix.

By varying the haplotype population in terms of number of haplotypes and pairwise distance while keeping the coverage constant, we generate five additional haplotype populations (population 2-6 as illustrated in Figure 5C). Across all populations, we again observe perfect precision of 1 for PredictHaplo. For populations 3 and 4, CliqueSNV has nearly perfect precision of 0.83*−*1. However, CliqueSNV is only able to detect haplotypes from the larger group of 20 haplotypes. Both CliqueSNV and PredictHaplo obtain their highest recall for populations 1 and 2 (Figure 4B), which are the two populations with only 10 haplotypes, and their lowest recall for populations 5 and 6 each with 55 haplotypes. This indicates that both tools are not able to appropriately deal with large haplotype populations. As before, CliqueSNV’s generally higher recall than PredictHaplo’s, is due to CliqueSNV predicting a larger amount of haplo-types than PredictHaplo. In all simulated populations, we observe that PredictHaplo predicts a single haplotype per cluster while CliqueSNV finds, if any, always multiple ones per cluster (Figure 4B). HaploClique and HaploConduct remain at a recall and precision of 0.

**Fig. 5:**
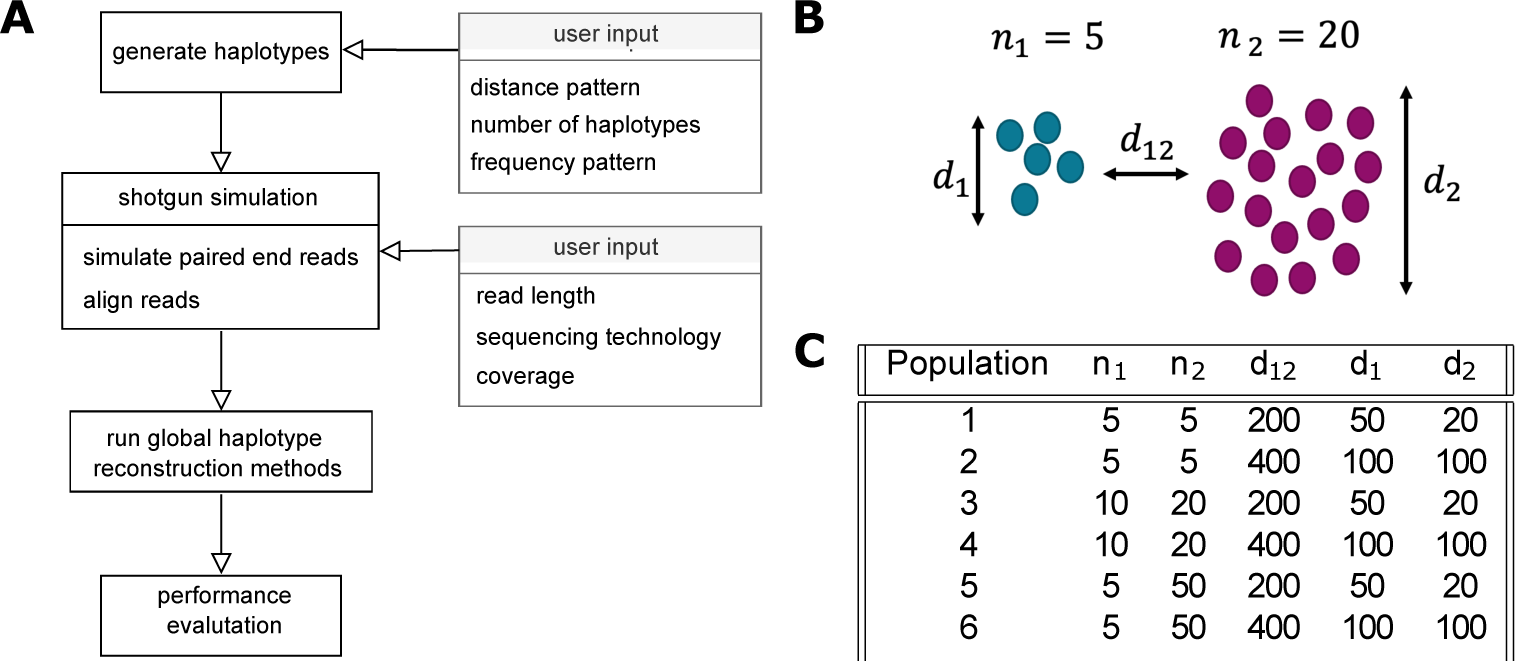
**A)** Workflow for the performance evaluation of global haplotype reconstruction methods: 1. Generation of haplotype population based on user input, 2. Simulation of paired-end Illumina sequencing reads, 3. Run global haplotype reconstruction methods, 4. Performance evaluation. **B)** Generation of distance based haplotype populations: *n*_1_: number of haplotypes in group one; *n*_2_: number of haplotypes in group two; *d*_12_: average pairwise distance between group one and two; *d*_1_: average pairwise sequence distance within group one; *d*_2_: average pairwise sequence distance within group two. **C)** Haplotype population parameter settings for the second synthetic dataset with constant coverage of 1000, and genome of length 10000.

Next, we used the experimental HIV-5 strain mixture [15] to evaluated the methods on a real sequencing data. We observe that precision and recall remain in the range of 0.2 *−* 0.4 for PredictHaplo. CliqueSNV and HaploConduct remain at 0 for precision and recall. As before, PredictHaplo’s and CliqueSNV’s reconstructions cover nearly the whole genome while HaploConduct reaches less than a fifth (Figure 4C).

In summary, our benchmark studies demonstrates that CliqueSNV exhibits the shortest runtime and delivers the highest recall performance for the simulated samples, whereas PredictHaplo exhibits superior precision for the same samples. This can mostly be explained by CliqueSNV typically recovering a larger amount of haplotypes than PredictHaplo. PredictHaplo was better able to reconstruct global haplotypes with the real data set both in terms of precision and recall. Overall, the results of our benchmark study indicate that the performance of all methods is diverse and highlights the need of continuous benchmarking as new methods are developed.

The benchmarking study can be effortlessly reproduced due to its adherence to Snakemake’s guidelines. It can be easily customized for different scenarios by integrating a novel data generation script. Moreover, incorporating new methods into the study merely requires adding a short script to execute those methods. Thus, our benchmarking study itself aligns with sustainable data processing practices.

## 4 Comparison to other workflows

We compare V-pipe 3.0 to other relevant viral bioinformatics pipelines for within-sample diversity estimation, focusing on functionalities and sustainability (Table 2). The compared pipelines include nf-core/viralrecon [10], HAPHPIPE [11] and ViralFlow [9]. These pipelines are all open source, actively maintained, and provide within-sample diversity estimates for Illumina sequencing reads. Active maintenance is crucial in this rapidly evolving field as even frequently used methods are still in continuous development and contain bugs for corner cases that only become evident with the rise of massive data sets in recent years. During the SARS-CoV-2 pandemic many processing pipelines have been developed, however the vast majority of those are specific to SARS-CoV-2, tailored to the ARTIC protocol [55] combined with Illumina sequencing, and only aim to produce consensus sequences. Since SARS-CoV-2 has limited genetic diversity and a well-known reference sequence, these pipelines cannot be easily adapted for the general case.

**Table 2:** Comparison in terms of sustainability and functionalities of viral bioinformatic workflows for within-sample diversity estimation.

|  | V-pipe 3.0 | ViralFlow | nf-core/viralrecon | HAPHPIPE |
| --- | --- | --- | --- | --- |
| **Reproducibility** |  |  |  |  |
| Automatic installation of all software dependencies | ✓ | ✓ | ✓ | ✗ |
| Container Services (e.g. Docker) | ✓ | ✓ | ✓ | ✗ |
| Automatic pipeline installation tests | ✓ | ✗ | ✓ | ✗ |
| Automatic pipeline execution tests on experimental samples | ✓ | ✗ | ✓ | ✗ |
| **Scalability** |  |  |  |  |
| Dynamic cluster resource allocation | ✓ | ✓ | ✓ | ✗ |
| **Adaptability** |  |  |  |  |
| Applicable for general viruses | ✓ | ✗ | ✓ | ✓ |
| Modular execution | ✓ | ✗ | ✓ | ✓ |
| Development: feature adding | ✓ | ✗ | ✓ | ✗ |
| **Transparency** |  |  |  |  |
| Open source | ✓ | ✓ | ✓ | ✓ |
| Readability: Standard pipeline code structure | ✓ | ✗ | ✓ | ✗ |
| Documentation | ✓ | ✓ | ✓ | ✓ |
| Tutorials and examples | ✓ | ✓ | ✗ | ✓ |
| **Functionalities** |  |  |  |  |
| De novo assembly | ✓ | ✗ | ✓ | ✓ |
| Read alignment | ✓ | ✓ | ✓ | ✓ |
| Consensus sequence generation | ✓ | ✓ | ✓ | ✓ |
| Mutation calling | ✓ | ✓ | ✓ | ✓ |
| Local haplotype reconstruction | ✓ | ✗ | ✗ | ✗ |
| Global haplotype reconstruction | ✓ | ✗ | ✗ | ✓ |
| SARS-CoV-2 wastewater surveillance | ✓ | ✓ | ✗ | ✗ |

The pipeline ViralFlow, however, also provides variant calling for Illumina sequencing reads and downstream analysis for SARS-CoV-2 lineage assignment. In terms of functionality, all data processing pipelines enable *de novo* assembly, except for ViralFlow. HAPHPIPE and nf-core/viralrecon use SPAdes [56] for this purpose, while V-pipe 3.0 utilizes Vicuna [20]. For read alignment, consensus sequence generation, and single nucleotide variant calling, each pipeline offers different combinations of tools and methods. For instance, both ViralFlow and nf-core/viralrecon provide the option to use iVar’s variant calling and consensus sequence generation. HAPHPIPE uses GATK for variant calling, and V-pipe 3.0 integrates two mutation callers: LoFreq and ShoRAH, which also provides local haplotypes. V-Pipe 3.0 stands out with its integrated benchmarking framework (Section 7.1). This framework allows for simulation of sequencing reads from flexible haplotype populations and performance evaluation of various methods. In contrast, [57] presented a benchmarking workflow for a global haplotype caller that is not easily adaptable due to hard-coded simulation parameters in bash-scripts.

Apart from its functionalities, sustainability is an essential factor for data analysis of enduring impact. V-pipe 3.0, ViralFlow, and nf-core/viralrecon ensure reproducibility and portability by providing software dependency definitions, automatically installing all necessary dependencies upon pipeline installation or execution. HAPHPIPE, on the other hand, requires manual installation of some software dependencies. In addition, V-pipe 3.0, ViralFlow, and nf-core/viralrecon offer container services like Docker, ensuring full pipeline portability and reproducibility. All four pipelines are transparent and open source, utilizing publicly available tools and methods. They provide documentation for installation and execution. In addition, HAPHPIPE and V-pipe 3.0 offer tutorials and examples to aid users in applying the pipelines to their data. Both nf-core/viralrecon and V-pipe 3.0 have code structures that conform to recommended standards for Nextflow and Snakemake workflows, ensuring code readability for external users, which makes adding new features straightforward. The other work-flows follow more custom code structures, making it challenging to add new features or modify the workflow, thus limiting their adaptability.

Overall, with their portability, automatic tests and gold standard code structure, the workflows nf-core/viralrecon and V-pipe 3.0 can provide sustainable data processing and analysis. While HAPHPIPE and V-pipe 3.0 provide the broadest range of functionalities with additional options for downstream analysis like phylogenetic tree building, analysis of co-occurrence of mutations on amplicons (COJAC), or kernel-based deconvolution for time-series frequency curves of variants (LolliPop).

Further, V-pipe 3.0 integrates the largest selection of tools for each processing step to ensure suitable processing for different samples. For example, for alignment V-pipe 3.0 supports BWA MEM, Bowtie 2, ngshmmalgin and minimap2.

## 5 Discussion

We have presented V-pipe 3.0, a sustainable data analysis pipeline designed for analyzing next-generation sequencing data of short viral genomes. In particular, we describe how we designed it to be reproducible by following Snakemake’s best-practice guidelines, adaptable by implementing virus-specific configuration files which can be quickly exchanged, and transparent by providing automatically tested usage examples, which are available online. We demonstrate the effectiveness and utility of these developments by highlighting its application to two large-scale projects, where V-pipe 3.0 was used in a production setting to process thousands of samples over multiple years.

One of V-pipe 3.0’s core functionalities is the estimation of viral diversity from NGS data. To address this challenge, we have developed a versatile benchmarking module that facilitates the continuous assessment of the performance and limitations of existing diversity estimation methods. As this field is still quickly advancing, continuous benchmarking of new and established methods is needed. For this purpose we focus on making the addition of new tools and test data sets to the workflow as straightforward as possible. Adding new methods is as easy as writing a single script which defines how to execute the tool and how to install it. New data sources can be either synthetic or derived from real experimental samples. In the synthetic case, different haplotype evolution modeling assumptions can be specified in a flexible way. Real data sources can be automatically downloaded and pre-processed as part of the workflow.

Given the mixed performance observed in our benchmark study for global haplo-type reconstruction, it is evident that the current methods may not satisfy the demands of downstream applications. The issues with performance can be attributed not only to the limitations of inference methods but also to the complex population structures inherent to viruses. Consequently, the practical application of global haplotype reconstruction is heavily constrained by these poor performing and often non-scalable methods, and would require improved scalable methods that explicitly account for the uncertainty of the results.

When comparing V-pipe 3.0 to other pipelines with similar purposes we found that, apart from V-pipe 3.0, only nf-core/viralrecon provides sustainable data processing taking into account reproducibility, portability, adaptability and transparency by following Nextflow’s best-practice guidelines. V-pipe 3.0 sets itself apart from the other pipelines by offering a broader range of integrated tools and functionalities, supported by thorough documentation and tutorials that address various application settings.

## 6 Conclusions

In summary, we have developed V-pipe 3.0 a sustainable data analysis pipeline for within-sample diversity estimation that can be easily applied to large numbers of samples by other researchers while keeping its execution robust and its workflow structure open to modifications. We have created a benchmarking module for one of V-pipe 3.0’s core functionalities which can be continuously updated when new methods and data sets appear. By continuing our close contact and exchange with users through our mailing list, active GitHub discussions and workshops, we will further expand V-pipe 3.0 to support different kinds of sequencing data, make it more robust to unpredictable failure points in cluster environments and further improve interoperability with data providers and consumers.

## 7 Methods

In the following, we introduce V-pipe 3.0’s benchmarking module and its application to the global haplotype reconstruction benchmarking study in detail.

### 7.1 Benchmarking module

V-pipe 3.0’s benchmarking module allows the benchmarking of global haplotype reconstruction methods on real and simulated data. For simulated data the workflow consists of four steps: generation of haplotype populations, shotgun read simulation, methods execution and performance evaluation (Figure 5A). In the case of real data, the first two steps are replaced by a data downloading and alignment step.

#### Generation of synthetic data sets

The synthetic data sets are generated in two steps. First, viral haplotype populations are generated. In the second steps, reads are simulated (Figure 5A). If no reference sequence is provided by the user, it is generated by drawing bases uniformly at random for each position based on the user-provided genome length.

We integrated two options for the viral haplotype population generation based on user-specified mutation rates or pairwise distances. Incorporating new methods involves the addition of a new script to the module, which generates haplotypes in fasta format as output. In the case of haplotype generation based on mutation rates, substitutions, deletions and insertions are randomly introduced into the master sequence based on the user-specified rates *µ*. The frequency composition of those haplotypes in the population is derived from haplotype frequencies *f* = (*f*_1_*,…, f_K_*) provided by the user. These simulation settings allow testing the reconstruction limits of the different viral diversity estimation methods.

In the case of haplotype generation by pairwise distances, we simulate hierarchical relationships among the haplotypes by generating two groups of closely related haplotypes that share a common ancestor (Figure 5B). First, using the user-specified between-group pairwise distance *d*_12_ two haplotypes are generated from the reference sequence. Second, for each haplotype, child-haplotypes are generated by introducing mutations based on the respective within-group pairwise distance (*d*_1_ and *d*_2_ respectively) and group size (*n*_1_ and *n*_2_ respectively). The frequency distribution of the generated haplotypes is obtained from a geometric series with a given ratio (default: 0.75), this results in a few high-frequency and many low-frequency haplotypes being present. Additionally the frequency distribution can also be drawn from a Dirichlet distribution with user-povided concentration parameters *α_i_*.

Given a user-specified per-position coverage and read length, paired-end reads are simulated in shotgun-mode using the ART Illumina read simulator [58].

#### Integration of real data sets

In addition to synthetic data sets where the ground truth is known, real data sets are included in the benchmark. We test the global haplotype reconstruction methods on sequencing reads from the 5-virus-mix presented in [15]. It provides Illumina MiSeq reads for a mixture of five HIV-1 strains: HXB2, 89.6, JR-CSF, NL4-3 and YU-2 and thus gives an estimate of the ground truth which can be used for performance evaluation. The benchmark workflow is designed to make the addition of further real data sets easily possible.

#### Performance evaluation

To evaluate the performance of each method in the global haplotype reconstruction benchmark, we compute precision and recall for the recovery of ground truth global haplotypes for each method in each condition. To do so, we consider the ground truth set of haplotype sequences and the set of sequences produced by a method. For each predicted sequence, we check if there exists a ground truth sequence with a relative edit distance below a predefined threshold *γ*. We define the relative edit distance *ED_rel_* as

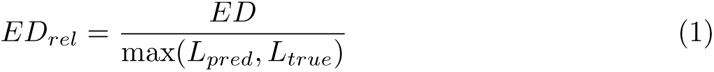

where *ED* is the edit distance between a predicted and ground truth haplotype which have lengths *L_pred_*and *L_true_* respectively. If *ED_rel_ < γ*, the predicted haplotype counts as a true positive, otherwise as a false positive. To compute the number of false negatives, we iterate over all ground truth sequences. We count a false negative if a ground truth sequence has no matching, i.e., relative edit distance below a certain threshold, predicted sequence. From this, we compute precision as *TP/*(*TP* + *FP*) and recall as *TP/*(*TP* + *FN*). We use *γ* = 0.01 as the relative edit distance threshold in the benchmark study.

Two-dimensional embeddings of haplotype sequences are generated by applying multidimensional scaling with precomputed edit distances between all sequences [59].

We use MetaQUAST to compute measures of assembly quality for the reconstructed haplotypes [60]. In particular, we compute the N50 score which, in this context, equals the length of the shortest haplotype, which together with all larger haplotypes, covers at least half the genome.

### 7.2 Global haplotype reconstruction benchmark study

We used the benchmarking module to benchmark global haplotype reconstruction methods.

#### Datasets

We generated two synthetic data sets applying the distance-based haplotype generation mode and used one real data set. In the first synthetic data set, we considered a genome of length 10000 with reads of length 200. We then generated two groups of haplotypes such that group one has size *n*_1_ = 5 and group two has size *n*_2_ = 5, the average pairwise sequence distance within group one is *d*_1_ = 50, the average pairwise sequence distance within group two is *d*_2_ = 20, and the average pairwise sequence distance between the two groups is *d*_12_ = 200. We varied the coverage between 500, 1000, 5000, 10000 in order to investigate how well the methods are able to recover low-frequency haplotypes as the coverage decreases. In the second synthetic data set, we considered a genome of length 10000 with reads of length 200 at a constant coverage of 1000. We then used the six haplotype population parameter settings as specified in Figure 5C in order to investigate how well the methods are able to recover different types of haplotype populations with different diversity levels. For the real data set, we used the 5-virus-mix which contains the HIV-1 strains HXB2, 89.6, JR-CSF, NL4-3 and YU-2 mixing in uniform proportions.

#### Global haplotype methods

We considered all methods discussed in [54] for which a Conda package is available. They are aBayesQR [61], CliqueSNV [62], HaploClique [31], HaploConduct [30], PEHaplo [63], PredictHaplo [29], QuasiRecomb [32], and RegressHaplo [64]. From the benchmark study we excluded aBayesQR because the program failed to parse the input sequencing reads, PEHaplo because it failed execution during the result assembly, QuasiRecomb as it terminated during startup and Regresshaplo, because not all dependencies of its Conda package were available.

The remaining tools are HaploConduct, HaploClique, PredictHaplo and CliqueSNV which are all reference-based global haplotype reconstruction methods. This means that they rely on the existence of a viral reference sequence which is similar to the haplotypes expected to occur. The input reads are then typically mapped against this reference sequence which makes reconstructing global haplotypes easier, because read positions relative to the genome are available, but also introduces a bias, as haplotypes which are dissimilar to the reference might not be captured. For the real data set, we had to exclude HaploClique for its excessive memory consumption.

## Declarations

### Ethics approval and consent to participate

Not applicable.

### Consent for publication

Not applicable.

### Availability of data and materials

V-pipe 3.0 is publicly available on GitHub [17]. All data and code for reproducing the benchmarking study is available on GitHub [65].

### Competing interests

The authors declare that they have no competing interests.

## Funding

LF was funded by European Union’s Horizon 2020 research and innovation program, under the Marie Sk-lodowska-Curie Actions Innovative Training Networks grant agreement no. 955974 (VIROINF).

## Authors’ contributions

LF, KPJ, IT and NB worked on the conceptualization and design of the pipeline. IT, KJP, LF, AAB, NBorg, PIB, MC, CC, AD, MD, DD, AJ, BL, MO and US were involved in implementing or adding new methods or tools. KJP conducted the benchmark study. CC, DD, IT, LdP, TS, MC, FS, NB, LF, and KPJ were involved in the analysis and processing of the SARS-CoV-2 clinical and wastewater samples. DD, IT, NB, KJP, LF were involved in the visualization of the results. KPJ and LF were writing the original draft. NB, LdP, TS, FS were involved in reviewing and editing of the manuscript. All authors read and approved the final manuscript.

## Supporting information

Detailed acknowledgements of the originating and submitting laboratories of the GISAID data.

## Acknowledgements

We gratefully acknowledge all data contributors, i.e., the Authors and their Originating laboratories responsible for obtaining the specimens, and their Submitting laboratories for generating the genetic sequence and metadata and sharing via the GISAID Initiative [66].

## Notes

### Competing Interest Statement

The authors have declared no competing interest.

