## Supplementary material for "V-pipe 3.0: a sustainable pipeline for within-sample viral genetic diversity estimation": Detailed acknowledgements of the originating and submitting laboratories of the GISAID data.

### SUPPLEMENTAL TABLE

**Data Availability**


| | |
|---|---|
| GISAID Identifier: | EPI_SET_231013cd |
| doi: | 10.55876/gis8.231013cd |


All genome sequences and associated metadata in this dataset are published in GISAID's EpiCoV database. To view the contributors of each individual sequence with details such as accession number, Virus name, Collection date, Originating Lab and Submitting Lab and the list of Authors, visit 10.55876/gis8.231013cd

**Data Snapshot**

- EPI_SET_231013cd is composed of 76,546 individual genome sequences.
- The collection dates range from 2020-03-04 to 2022-12-30;
- Data were collected in 1 countries and territories;
- All sequences in this dataset are compared relative to hCoV-19/Wuhan/WIV04/2019 (WIV04), the official reference sequence employed by GISAID (EPI_ISL_402124). Learn more at https://gisaid.org/WIV04.
